# Increased 5-HT_2A_ receptor signalling efficacy differentiates serotonergic psychedelics from non-psychedelics

**DOI:** 10.1101/2024.06.13.594677

**Authors:** Aurelija Ippolito, Sri Vasudevan, Shaun Hurley, Gary Gilmour, Frederick Westhorpe, Grant Churchill, Trevor Sharp

## Abstract

**Background and Purpose:** Serotonergic psychedelic drugs are under renewed investigation for the potential treatment of several psychiatric disorders. While all serotonergic psychedelics have 5-HT_2A_ receptor activity, the explanation for why some 5-HT_2A_ receptor agonists are not psychedelic is unknown. To address this question, we investigated the 5-HT_2A_ receptor signalling bias and efficacy of a panel of psychedelics and non-psychedelics.

**Experimental Approach:** G -coupled (Ca^2+^ and IP_1_) and β-arrestin2 signalling effects of eight chemically diverse psychedelics (psilocin, 5-MeO-DMT, LSD, mescaline, 25B-NBOMe and DOI) and non-psychedelics (lisuride and TBG) were characterised using SH-SY5Y cells expressing recombinant human 5-HT_2A_ receptors. Measurements of signalling efficacy and bias were derived from dose-responses curves for each agonist, compared to 5-HT. Follow-up experiments sought to confirm the generality of findings using rat C6 cells expressing endogenous 5-HT_2A_ receptors.

**Key Results:** In SH-SY5Y cells, all psychedelics were partial agonists at both 5-HT_2A_ receptor signalling pathways and none showed significant signalling bias. In comparison, in SH-SY5Y cells the non-psychedelics lisuride and TBG were not distinguishable from psychedelics in terms of biased agonist properties, but both exhibited the lowest 5-HT_2A_ receptor signalling efficacy of all drugs tested, a result confirmed in C6 cells.

**Conclusion and Implications:** In summary, all psychedelics tested were unbiased, partial 5-HT_2A_ receptor agonists. Importantly, the non-psychedelics lisuride and TBG were discriminated from psychedelics, not through biased signalling but rather by relatively low efficacy. Thus, 5-HT_2A_ receptor signalling efficacy and not bias provides a possible explanation for why some 5-HT_2A_ receptor agonists are not psychedelic.

## INTRODUCTION

Serotonergic psychedelics are in development for the treatment of psychiatric disorders ranging from major depression and anxiety to substance misuse disorder and anorexia^1,2^. Each of psilocybin, 5-methoxy-N,N-dimethyltryptamine (5-MeO-DMT) and lysergic acid diethylamide (LSD) have progressed to clinical trials of treatment-resistant depression and anxiety^3–5^. As a result, there is increasing interest in the molecular mechanisms by which such agents induce their characteristic psychedelic effects.

There is clear evidence that engagement at the 5-HT_2A_ receptor is central to the subjective effects of serotonergic psychedelics. Specifically, clinical PET imaging studies using the 5-HT_2A_ radioligand, [^11^C]Cimbi-36, report a strong correlation between 5-HT_2A_ receptor occupancy and the intensity of psychedelic effects following psilocybin administration^6,7^. Moreover, the subjective effects of both psilocybin and LSD in human volunteers were attenuated by the 5-HT_2A_ receptor antagonist ketanserin^8–12^. In preclinical studies, 5-HT_2A_ receptor agonist-induced head-twitches in mice are widely considered a surrogate marker of the psychedelic effects of these drugs in humans. Indeed, this head-twitch response is abolished by 5-HT_2A_ receptor knockout or selective antagonists^13,14^, and agonist potency in this model correlates with potency to induce psychedelic effects in humans^15^.

A recent interesting development is evidence of non-psychedelic 5-HT_2A_ receptor agonists. Thus, there are several reports of 5-HT_2A_ receptor agonists lacking the propensity to evoke head-twitches^16–18^. For example, in mice, administration of the high affinity 5-HT_2A_ receptor ligand tabernanthalog (TBG) did not induce head-twitches but was capable of evoking 5-HT_2A_ receptor-mediated effects in other in vivo models^18^. Similarly, lisuride, another high affinity 5-HT_2A_ receptor agonist, lacked effects on head-twitches in mice^16^. Although it is not yet known whether TBG is non-psychedelic when administered to humans, there are numerous clinical reports showing that lisuride lacks psychedelic effects^19–21^. The explanation for why some 5-HT_2A_ receptor agonists are psychedelic and not others, is unknown.

A common feature of G protein-coupled receptors is their capacity to signal through both G protein-dependent and β-arrestin2-dependent pathways^22–24^. Thus, the 5-HT_2A_ receptor has been shown to signal via G (to activate phospholipase C and increase inositol trisphosphate and intracellular Ca^2+^ ^7^) as well as β-arrestin2 and other pathways^25–28^. Divergence in the subjective effects of drugs with 5-HT_2A_ agonist activity could be driven by selective signalling through G_q_- or β-arrestin2-mediated pathways (biased agonism)^29^. As an example, biased agonism at μ-opioid receptors was initially thought to explain the preferential sedative versus analgesic effects of certain μ-opioid receptor agonists^30,31^. An alternative explanation for non-psychedelic 5-HT_2A_ receptor agonists is partial agonism; agonists with low of 5-HT_2A_ receptor efficacy may exhibit a more limited repertoire of behavioural effects^32–34^. Partial agonism at the benzodiazepine binding site of the GABA_A_ receptor was offered to account for the behaviourally selective effects of benzodiazepine agonists^35–37^. Moreover, partial agonism at μ-opioid receptors is the currently favoured alternative explanation for analgesia selective μ-opioid receptor agonists^30,31^.

Against this background, the current study investigated the 5-HT_2A_ receptor-mediated G_q_ or β-arrestin2 signalling properties of a panel psychedelics; the tryptamines psilocin (active metabolite of psilocybin) and 5-MeO-DMT, the ergoline LSD and the phenethylamines mescaline (3,4,5-trimethoxyphenethylamine), 4-bromo-2,5-dimethoxy-N-(2-methoxybenzyl)phenethylamine (25B-NBOMe) and 2,5-dimethoxy-4-iodoamphetamine (DOI). The signalling properties of these agents were compared with two non-psychedelics TBG and lisuride (also of the ergoline chemical class). The drugs were selected to be chemically diverse since receptor stabilization in a particular state might determine signalling bias^16,38–40^. Experiments utilised cell lines expressing human or rat 5-HT_2A_ receptors.

## METHODS

### Cell culture

SH-SY5Y neuroblastoma cells transfected with the human 5-HT_2A_ receptor^41^ were maintained in culture medium; Dulbecco’s Modified Eagle Medium (DMEM) containing 2 mM Glutamax, 10 % (v/v) fetal bovine serum (FBS), 100 I.U. µg^-1^ ml^-1^ penicillin/streptomycin, and 480 µg ml^-1^ G418 (to maintain transfection selection pressure). C6 glioma cells which endogenously express the rat 5-HT_2A_ receptor^42,43^ (ATCC CCL-107) were maintained in Ham’s F12 nutrient mix containing 2 mM Glutamax, 10 % (v/v) FBS and 100 I.U. µg^-1^ ml^-1^ penicillin/streptomycin. Cells were grown at 37°C in a humidified atmosphere of 95 % air and 5 % CO_2_.

### Assay of agonist-evoked cytosolic Ca

Cells were plated in 96-well black/clear bottom plates at a density of 40,000 (SH-SY5Y) or 60,000 (C6) cells/well, 48 h (SH-SY5Y) or 24 h (C6) before the day of experiment. The 10 % FBS culture medium was replaced with culture medium containing 10 % dialysed-FBS to avoid potential receptor desensitisation by 5-HT in the FBS.

On the day of experiment, the culture medium was aspirated and cells were washed twice with 200 µl Hanks’ Balanced Salt Solution (HBSS) containing calcium (HBSS-Ca^2+^). Next, 100 µl of assay buffer containing 4 µM Fluo-4-AM (Life Technologies), 0.02 % Pluronic F127 and 2.5 mM probenecid in HBSS-Ca^2+^ was added to each well and the plate was incubated at room temperature (RT) for 1 h to allow dye loading, followed by 37°C for 30 min to allow for intracellular esterase action. The assay buffer was aspirated and cells were washed twice with HBSS-Ca^2+^ before addition of 90 µl/well HBSS-Ca^2+^. Cells were allowed to equilibrate for 15 min at RT before fluorescence recordings at the same temperature.

Baseline fluorescence was measured on a plate reader (BMG Optima) from the plate bottom at 480/520nm excitation/emission every 5 s for 30 s prior to addition of one of; 10 µl agonist, 10 µl agonist/agonist combination or 10 µl agonist plus 10 µl antagonist (MDL-100,907 15 min before agonist addition), after which fluorescence was recorded for a further 2 min. In each well the final concentration of DMSO was 0.1 % (v/v).

### Assay of agonist-evoked inositol monophosphate (IP_1_) accumulation

SH-SY5Y cells were plated in 384-well white low-volume plates 24 h before the day of experiment at a density of 20,000 cells/well. As above, the 10 % FBS supplemented culture medium was replaced with 10 % dialysed-FBS. On the day of experiment, the medium was aspirated and 10 µl of buffer containing LiCl and 4 µl of agonist/antagonist was added to each well. The plate was incubated at 37°C for 1 h before addition to each well of 3 µl IP_1_-d2 and anti-IP_1_-d2 dissolved in lysis buffer (CisBio HTRF IP-One G_q_ Kit) and then incubated further at RT for 1 h. A calibration curve was run prior to commencing experiments (according to manufacturer’s instructions).

Fluorescence was measured on a plate reader (Tecan Infinite F1200) from the top of the plate at 620/340nm and 665/340nm excitation/emission. The final concentration of DMSO in each well was 0.1 % (v/v).

### Assay of agonist-evoked β-arrestin2 recruitment

SH-SY5Y cells were plated in 96-well black/clear-bottom plates at a density of 40,000 cells/well 48 h before the day of experiment. The 10 % FBS supplemented culture medium was replaced with 10 % dialysed-FBS. The following day, to each well was added 50 µl of a transfection mix containing 8 µl of β-arrestin2 sensor, 15 µl of human-5-HT_2A_ receptor, 3 µl GPCR kinase 2, 3 µl GPCR kinase 3 (all packaged in Mammalian Baculovirus vectors, Montana Molecular), 0.6 µl sodium butyrate and 21.4 µl media. The transfection mix was aspirated and cells washed twice with DPBS containing calcium (DPBS-Ca^2+^) before addition of 100 µl DPBS-Ca^2+^ to each well. The cells were allowed to equilibrate for 30 min at RT.

Baseline fluorescence was measured on a plate reader (BMG Omega) from the bottom of the plate at 485/520 excitation/emission every 15 s for 60 s prior to addition of 50 µl agonist, after which fluorescence was recorded for a further 20 min. The final concentration of DMSO in each well was at 0.1 % (v/v).

### Drugs

Psilocin (supplied by Compass Pathways), 5-MeO-DMT (Cambridge Bioscience), DOI (Cambridge Bioscience), mescaline (Cambridge Bioscience), LSD (Chiron), 25B-NBOMe (Chiron), lisuride (Bio-Techne), TBG (supplied by Compass Pathways) and MDL-100,907 (volinanserin; Bio-Techne), were dissolved in DMSO, and 5-HT-HCl (Enzo Life Sciences) was dissolved in deionised water, to obtain 10 mM stock solutions. On the day of experiment, working solutions were obtained by diluting drugs in HBSS-Ca^2+^ (Ca^2+^/IP_1_ assays) or DPBS-Ca^2+^ (β-arrestin2 assays).

### Data processing and statistical analysis

Statistical analyses were performed using GraphPad Prism software. For Ca^2+^ assays, baseline fluorescence was averaged and maximum fluorescence reached was expressed as a % of baseline (corrected for vehicle addition). For IP_1_ assays, data were converted to emission at 665/620 values which were then interpolated as intracellular IP_1_ concentrations using the IP_1_ calibration curve. For β-arrestin2 assays, fluorescence was averaged over the first 2.5 min and then measurements after agonist addition over the remaining 20 minutes were normalised to baseline. Steady states were then calculated using the ‘Baseline then rise to steady state time course’ curve fit on GraphPad Prism and this was used as an endpoint measurement of cumulative response.

Dose-response curves were generated using log[agonist] versus response (three parameter) curve fits (GraphPad Prism), which also provided potency and efficacy values. The dose response for each 5-HT_2A_ receptor agonist was normalised to the response to 10 µM 5-HT and each point was expressed as mean ± SEM value of at least two independent experiments carried out in duplicate.

The relative activity of agonists in different assays were calculated using the method of Kenakin et al^44^. Specifically, log(E_max_/EC_50_) values were calculated for each agonist in each pathway using E_max_ and EC_50_ values derived by averaging these parameters across replicates for each assay, and then compared to the reference agonist 5-HT by calculation of Δlog(E_max_/EC_50_) (log(E_max_/EC_50_)_agonist_ – log(E_max_/EC_50_)_5-HT_).

## RESULTS

### Effect of 5-HT_2A_ receptor agonists on cytosolic Ca^2+^ in SH-SY5Y cells

Initial experiments determined the effect of the selected psychedelic and non-psychedelic 5-HT_2A_ receptor agonists on G_q_ signalling activity via measurement of cytosolic Ca^2+^ increase in SH-SY5Y cells expressing recombinant human 5-HT_2A_ receptors. All drugs tested elicited dose-dependent increases in cytosolic Ca^2+^ (Fig. 1). The psychedelics tested had variable potencies with 25B-NBOMe and LSD being the most potent and mescaline the least potent (Fig. 1, Table 1).

**Figure 1.**
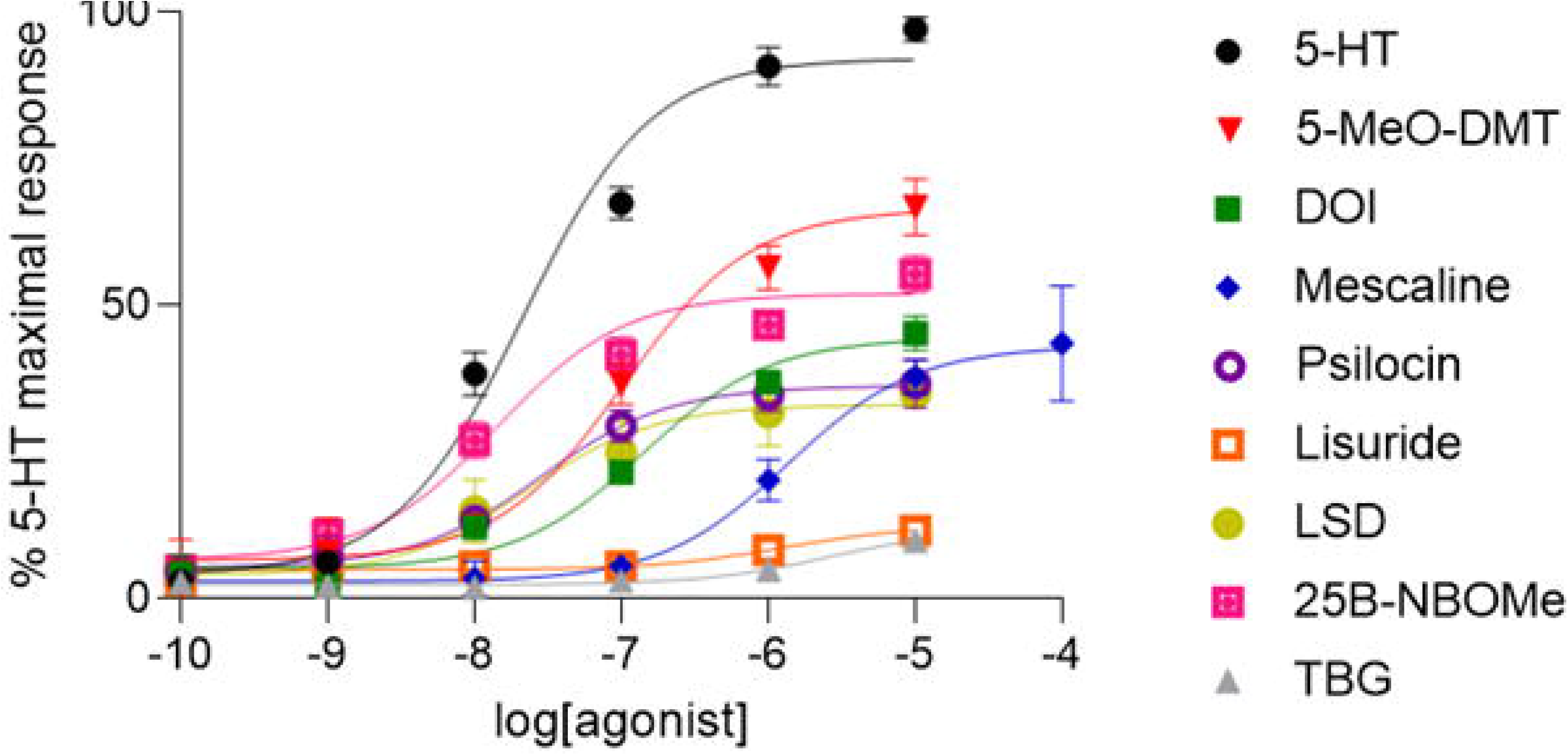
Effect of psychedelic and non-psychedelic drugs on cytosolic Ca^2+^ in SH-SY5Y cells expressing the human 5-HT_2A_ receptor. Each point is the mean ± SEM value of triplicates in two independent experiments. Responses are relative to 10 μM 5-HT.

**Table 1.** Signalling parameters of psychedelic and non-psychedelic drugs at the human 5-HT_2A_ receptor. Data were derived from dose-response curves for Ca, IP_1_ and β-arrestin2 readouts in SH-SY5Y cells expressing the human 5-HT_2A_ receptor. E_max_, pEC_50_ and Δlog(E_max_/EC_50_) values are the mean ± SEM of triplicates in two independent experiments. E_max_ and Δlog(E_max_/EC_50_) values are relative to 10 μM 5-HT. Non-psychedelic drugs are denoted by asterisk.

| Drug | Ca<sup>2+</sup> | | IP<sub>1</sub> | | β-arrestin2 | | Ca<sup>2+</sup> | IP<sub>1</sub> | β-arrestin2 |
| --- | --- | --- | --- | --- | --- | --- | --- | --- | --- |
| | E<sub>max</sub> | pEC<sub>50</sub> | E<sub>max</sub> | pEC<sub>50</sub> | E<sub>max</sub> | pEC<sub>50</sub> | $\Delta\log(E_{\max}/EC_{50})$ | $\Delta\log(E_{\max}/EC_{50})$ | $\Delta\log(E_{\max}/EC_{50})$ |
| **5-HT** | 1.00 ± 0.031 | 7.67 ± 0.10 | 1.00 ± 0.049 | 7.46 ± 0.17 | 1.00 ± 0.018 | 7.35 ± 0.063 | 0.00 | 0.00 | 0.00 |
| **5-MeO-DMT** | 0.73 ± 0.033 | 6.99 ± 0.18 | 0.93 ± 0.027 | 7.97 ± 0.10 | 0.70 ± 0.020 | 7.62 ± 0.095 | -0.82 ± 0.21 | 0.48 ± 0.18 | 0.10 ± 0.11 |
| **DOI** | 0.49 ± 0.019 | 6.85 ± 0.13 | 1.12 ± 0.025 | 7.48 ± 0.08 | 0.73 ± 0.027 | 8.12 ± 0.14 | -1.12 ± 0.17 | 0.06 ± 0.17 | 0.63 ± 0.16 |
| **Mescaline** | 0.47 ± 0.031 | 5.87 ± 0.18 | 0.81 ± 0.050 | 6.14 ± 0.18 | 0.64 ± 0.059 | 6.25 ± 0.26 | -2.12 ± 0.21 | -1.41 ± 0.20 | -1.30 ± 0.27 |
| **Psilocin** | 0.40 ± 0.019 | 7.55 ± 0.19 | 0.80 ± 0.030 | 7.40 ± 0.13 | 0.28 ± 0.019 | 8.19 ± 0.21 | -0.52 ± 0.22 | -0.16 ± 0.19 | 0.27 ± 0.22 |
| **Lisuride\*** | 0.14 ± 0.014 | 5.91 ± 0.33 | 0.48 ± 0.031 | 9.11 ± 0.26 | 0.21 ± 0.044 | 6.06 ± 0.43 | -2.62 ± 0.35 | 1.33 ± 0.24 | -1.99 ± 0.44 |
| **LSD** | 0.36 ± 0.027 | 7.64 ± 0.29 | 0.78 ± 0.039 | 9.14 ± 0.26 | 0.44 ± 0.019 | 8.04 ± 0.13 | -0.46 ± 0.31 | 1.57 ± 0.24 | 0.33 ± 0.15 |
| **25B-NBOMe** | 0.57 ± 0.021 | 7.84 ± 0.15 | 0.75 ± 0.040 | 8.95 ± 0.28 | 0.88 ± 0.017 | 8.20 ± 0.079 | -0.06 ± 0.18 | 1.37 ± 0.25 | 0.79 ± 0.10 |
| **TBG\*** | 0.13 ± 0.033 | 5.59 ± 0.52 | 0.35 ± 0.064 | 5.29 ± 0.29 | 0.19 ± 0.026 | 6.35 ± 0.42 | -2.96 ± 0.54 | -2.63 ± 0.26 | -1.74 ± 0.43 |

In terms of efficacy, all psychedelics had lower efficacy than 5-HT (Fig. 1), with psilocin and LSD being the least efficacious. Interestingly, the two non-psychedelics, lisuride and TBG, displayed the lowest efficacy of all drugs tested (Fig. 1, Table 1).

The Ca^2+^ responses of SH-SY5Y cells to both psychedelic and non-psychedelic drugs (10 µM) were abolished by pre-treatment with MDL-100,907 (1 µM), confirming the role of 5-HT_2A_ receptor activation in this G_q_ signalling activity (Suppl. Fig. 1A).

### Effect of 5-HT_2A_ receptor agonists on IP_1_ accumulation in SH-SY5Y cells

Next, the effect of the psychedelics and non-psychedelics on G_q_ signalling activity was determined upstream of the Ca^2+^ response by measuring of accumulation of intracellular IP_1_ in the SH-SY5Y cells. As with cytosolic Ca^2+^, all drugs tested elicited dose-dependent increases in IP_1_ (Fig. 2). The rank order of potency for IP_1_ accumulation varied across the psychedelics but was similar to the Ca^2+^ response, with 25B-NBOMe and LSD being the most potent psychedelics and mescaline the least potent (Fig. 2, Table 1).

**Figure 2.**
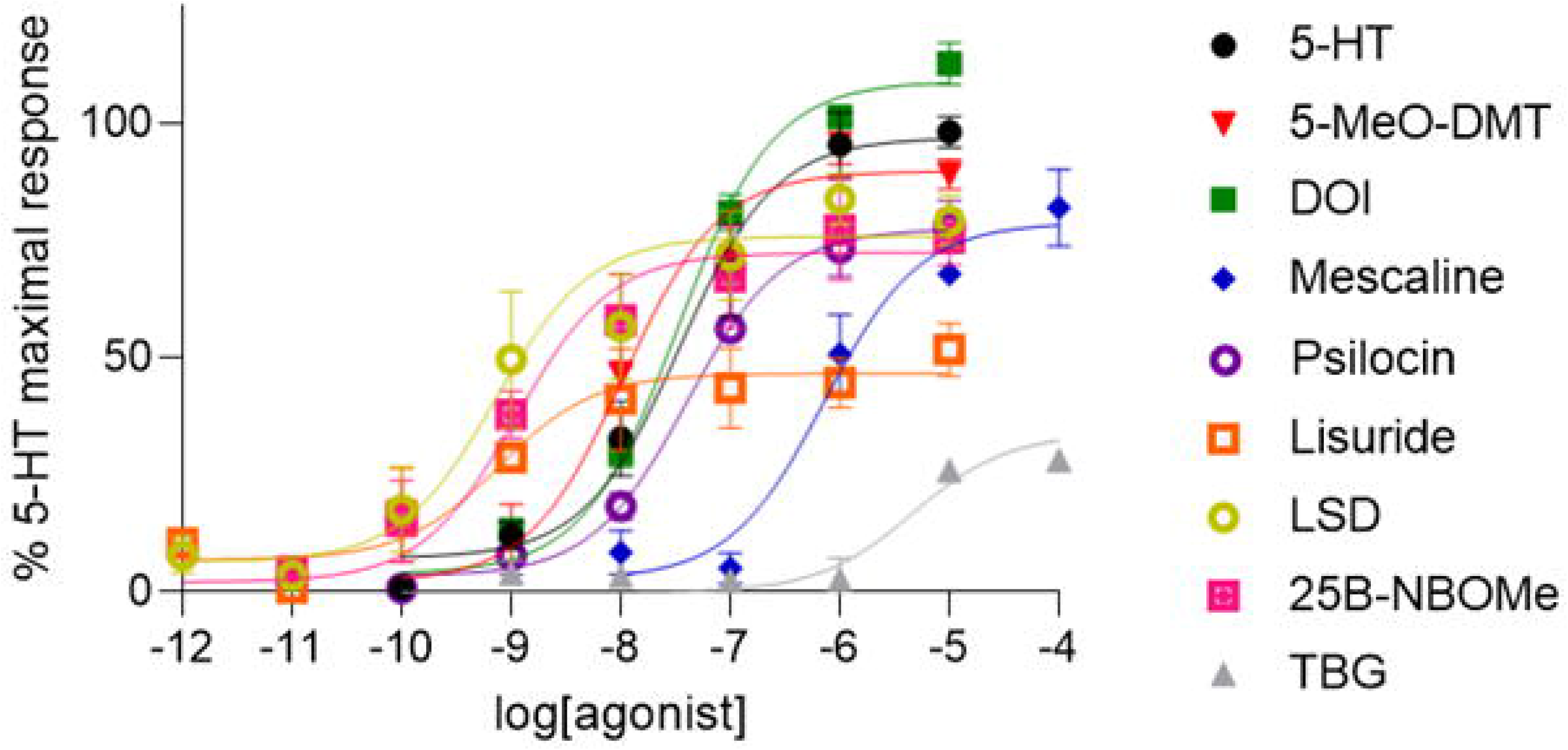
Effect of psychedelic and non-psychedelic drugs on IP_1_ accumulation in SH-SY5Y cells expressing the human 5-HT_2A_ receptor. Each point is the mean ± SEM value of triplicates in two independent experiments. Responses are relative to 10 μM 5-HT.

In the IP_1_ assay, most drugs were less efficacious than 5-HT, except 5-MeO-DMT and DOI. Interestingly, as noted for the Ca^2+^ assay, the non-psychedelics lisuride and TBG displayed the lowest efficacy of all drugs tested (Fig. 2, Table 1).

### Effect of 5-HT_2A_ receptor agonists on β-arrestin2 recruitment in SH-SY5Y cells

Next, experiments measured the effects of psychedelic and non-psychedelic drugs on 5-HT_2A_ receptor signalling via β-arrestin2 in the SH-SY5Y cells. As with the assays of G_q_ signalling, all drugs tested elicited dose-dependent increases in β-arrestin2 recruitment (Fig. 3). Moreover, 25B-NBOMe and LSD were amongst the most potent psychedelics tested, and mescaline amongst the least potent (Fig. 3, Table 1).

**Figure 3.**
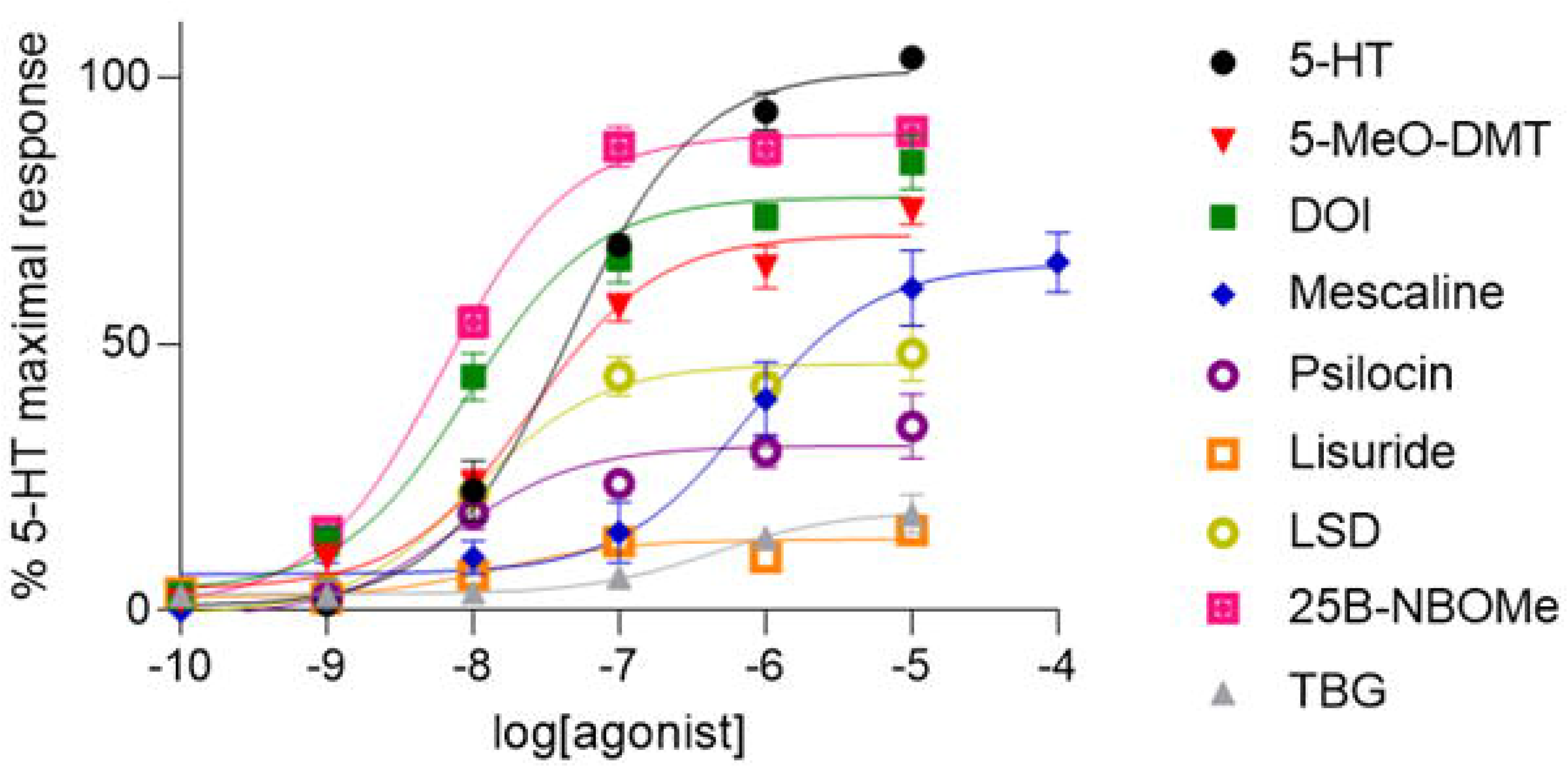
Effect of psychedelic and non-psychedelic drugs on β-arrestin2 recruitment in SH-SY5Y cells expressing the human 5-HT_2A_ receptor. Each point is the mean ± SEM value of triplicates in two independent experiments. Responses are relative to 10 μM 5-HT.

As observed in the Ca^2+^ assay, a feature of the β-arrestin2 assay was that drugs were less efficacious than 5-HT (Fig. 3, Table 1), in this case with psilocin being the least efficacious psychedelic. Moreover, of the drugs tested the non-psychedelics lisuride and TBG were ranked lowest in terms of efficacy.

The β-arrestin2 response to both psychedelics and non-psychedelics (10 µM) was abolished by pre-treatment with MDL-100,907 (1 µM), confirming that the β-arrestin2 signalling was 5-HT_2A_ receptor mediated (Suppl. Fig. 1B).

### Biased agonist properties of psychedelic and non-psychedelic agents

With agonist activity data generated from the IP_1_ and β-arrestin2 assays, ligand bias in 5-HT_2A_ receptor-mediated signalling was next assessed. For each assay, to cancel out system-specific differences such as downstream signalling amplification, agonist potencies and maximal efficacies were converted to Δlog(E_max_/EC_50_) values with 5-HT being the reference agonist. A scatter plot of Δlog(E_max_/EC_50_) values then allowed comparison of the activity of each agonist in these two pathways^44^ (Fig. 4). In this plot, agonists falling close to the line of unity (i.e. possessing similar Δlog(E_max_/EC_50_) values in each assay) have low bias, whereas agonists that deviate from the line of unity (i.e. possessing different Δlog(E_max_/EC_50_) values in each assay) display bias.

**Figure 4.**
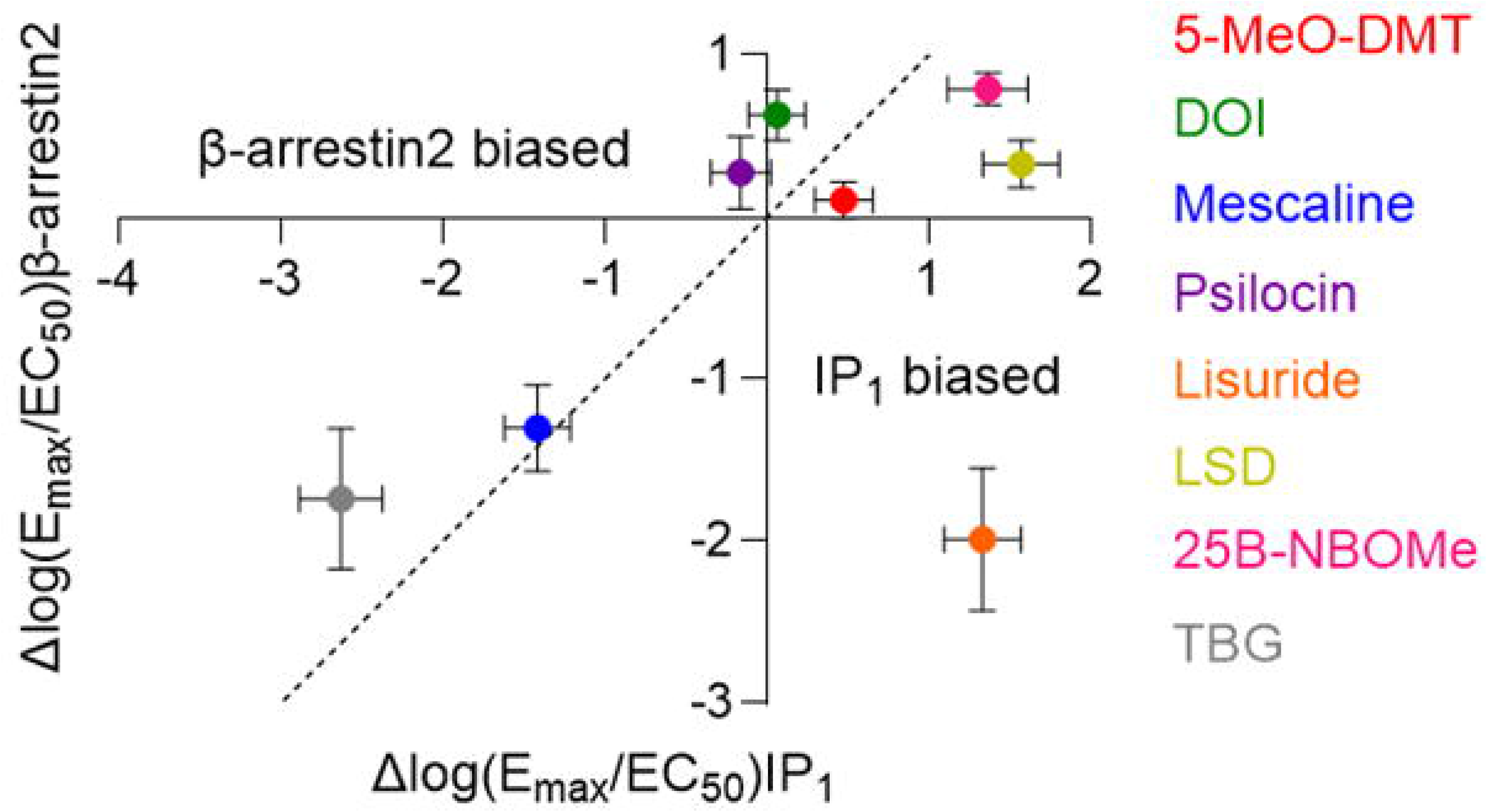
Scatter plot comparing the activity (Δlog(E_max_/EC_50_) values) of psychedelic and non-psychedelic drugs on 5-HT_2A_ receptor-mediated IP_1_ and β-arrestin2 signalling pathways in SH-SY5Y cells. In this plot, the more positive the x or y value, the greater activity in a particular pathway. The further a drug deviates from the line of unity, the more biased the agonist. Each point represents a mean ± SEM value.

There was a generally linear relationship between the IP_1_ versus β-arrestin2 signalling activity of the different agonists tested (Fig. 4). Of the psychedelic drugs, all were unbiased with the exception of LSD which showed a modest bias towards IP_1_ signalling versus β-arrestin2 signalling (Fig. 4). In comparison, one of the two non-psychedelics tested, lisuride, showed a bias towards IP_1_ versus β-arrestin2 signalling but TBG showed no signalling bias (Fig. 4).

Thus, comparison of agonist activity at IP_1_ versus β-arrestin2 signalling failed to distinguish between psychedelic and non-psychedelic 5-HT_2A_ receptor agonists. Overall, there was no clear pattern between signalling bias and chemical structure although it is noteworthy that the two ergolines LSD and lisuride showed bias towards IP_1_ signalling, albeit to varying degrees.

As with the plot of IP_1_ versus β-arrestin2 signalling, a scatter plot of Δlog(E_max_/EC_50_) values to compare agonist activity in the Ca^2+^ and β-arrestin2 signalling pathways did not discriminate between the psychedelic and non-psychedelic agonists (Suppl. Fig. 2A).

Finally, a scatter plot of Δlog(E /EC) values obtained from the Ca^2+^ and IP_1_ assays allowed comparison of agonist activity of what should be the same G_q_ signalling pathway. This revealed a generally linear relationship between the activity of the different agonists in the two pathways, although it was notable that LSD and lisuride showed increased activity in the IP_1_ assay compared to in the Ca^2+^ assay (Suppl. Fig. 2B).

Overall, the different scatter plots revealed, importantly, that signalling bias pattern did not distinguish between psychedelic and non-psychedelic agents.

### Effect of 5-HT_2A_ receptor agonists on cytosolic Ca^2+^ in C6 cells

Results obtained from the SH-SY5Y cells suggested that psychedelics could be distinguished from non-psychedelics on the basis of the latter having a very low efficacy at 5-HT_2A_ receptors and not differences in biased signalling activity. To test this observation further, the effect of drugs on G_q_ activity was examined in a different cell model, specifically C6 glioma cells endogenously expressing the rat 5-HT_2A_ receptor.

These experiments revealed that, as observed in SH-SY5Y cells, all psychedelics caused a dose-related increase in cytosolic Ca^2+^ in C6 cells (Fig. 5) with lower efficacy than 5-HT. Furthermore, the Ca^2+^ response to the psychedelics (10 µM) was abolished by pre-treatment with MDL-100,907 (1 µM), confirming 5-HT_2A_ receptor involvement (Suppl. Fig. 3B). The relative potency of the psychedelics in the C6 cells was similar to that observed in SH-SY5Y cells, with 25B-NBOMe and LSD being the most potent and mescaline being the least potent (Fig. 5; Table 2).

**Figure 5.**
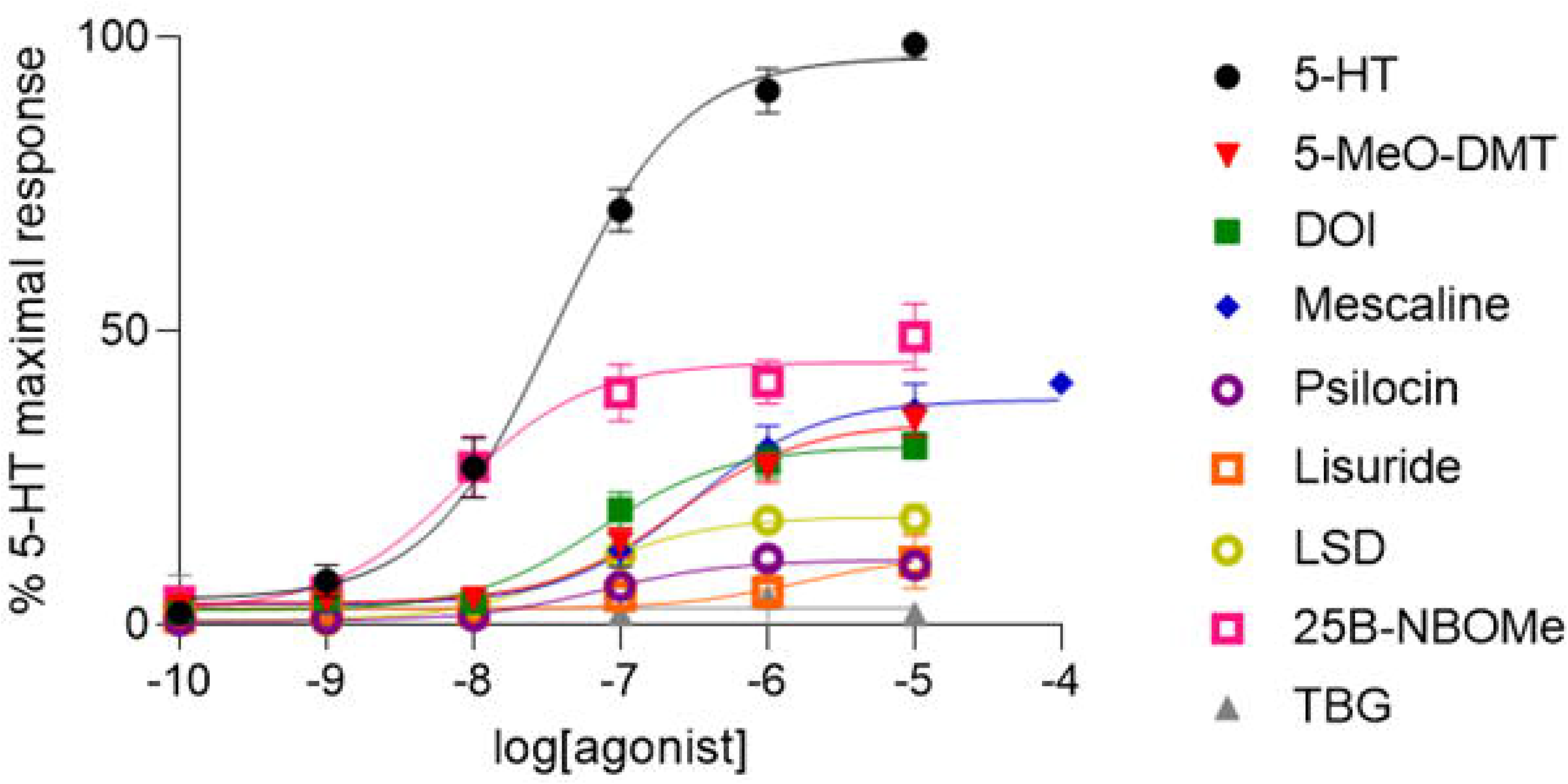
Effect of psychedelic and non-psychedelic drugs on cytosolic Ca^2+^ in C6 cells expressing the rat 5-HT_2A_ receptor. Each point is the mean ± SEM value of triplicates in two independent experiments. Responses are relative to 10 μM 5-HT.

**Table 2.**
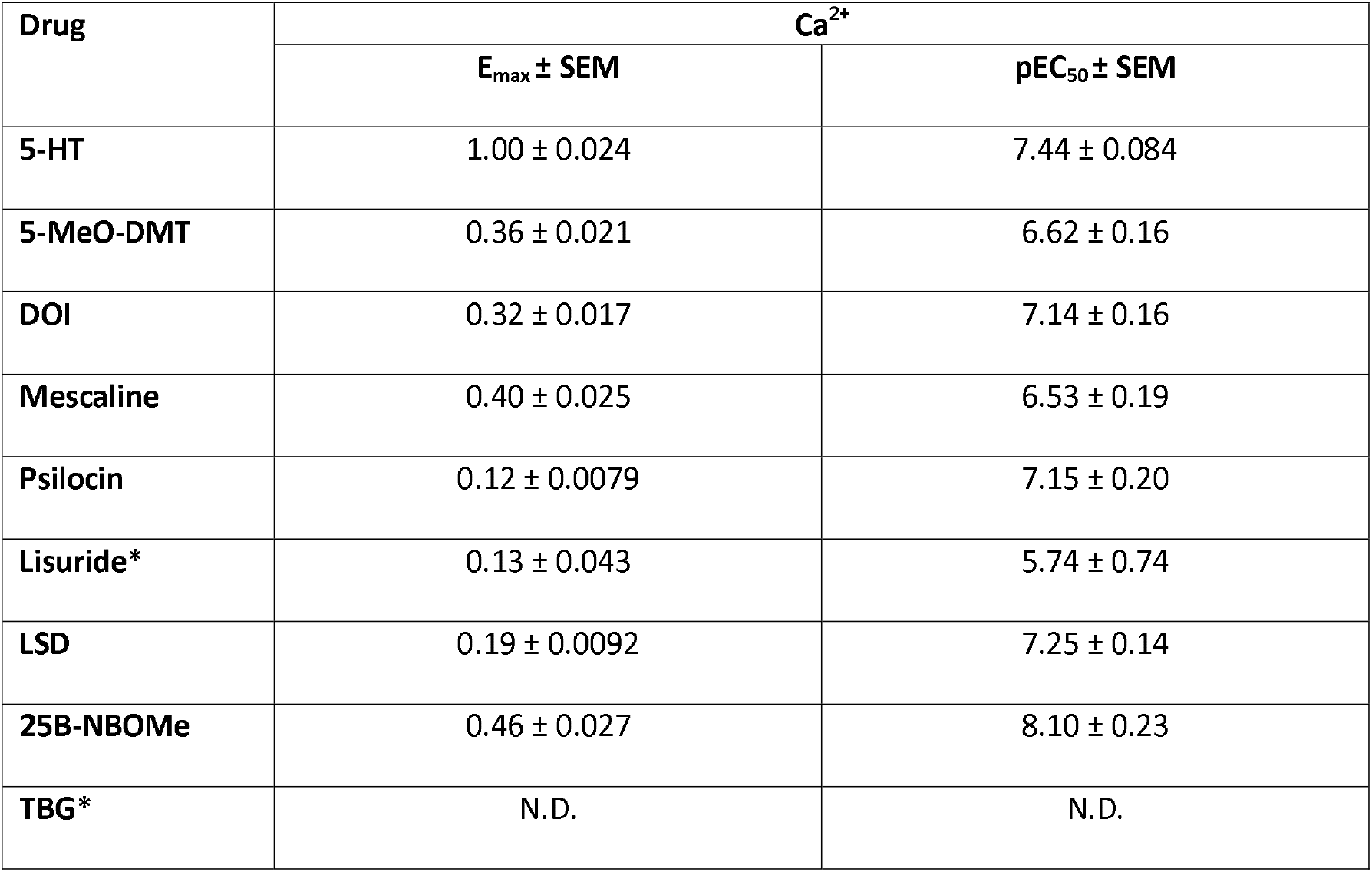
Potency and efficacy of psychedelic and non-psychedelic drugs at the rat 5-HT_2A_ receptor. Data were derived from dose-response curves for Ca readout in C6 cells expressing the rat 5-HT_2A_ receptor. E_max_ and pEC_50_ values are the mean ± SEM of triplicates in two independent experiments. E_max_ values are relative to 10 μM 5-HT. Non-psychedelic drugs are denoted by asterisk. N.D. – not detectable.

A Δlog(E /EC) scatter plot of Ca^2+^ responses emphasised the similarity in agonist activity in the C6 and SH-SY5Y cells (Suppl. Fig. 3A). However, it is noteworthy that mescaline was more active in C6 cells, and psilocin was more active in SH-SY5Y cells, potentially highlighting differences in the activity of these psychedelics at the rat versus human 5-HT_2A_ receptors.

Importantly, and also in keeping with results from the SH-SY5Y cells, the non-psychedelic lisuride had relatively low efficacy in the C6 cells and TBG had no measurable efficacy at the concentrations tested (Fig. 5; Table 2). Given their low efficacy both lisuride and TBG were run in combination with 5-HT, and Schild plots were constructed to confirm partial agonist properties. As expected of a partial agonist, the presence of both lisuride and TBG increased the response of low 5-HT concentrations and reduced the response of higher 5-HT concentrations (Fig. 6). Schild plots for the psychedelic agonists also confirmed the partial agonist properties of these drugs (Suppl. Table 1).

**Figure 6.**
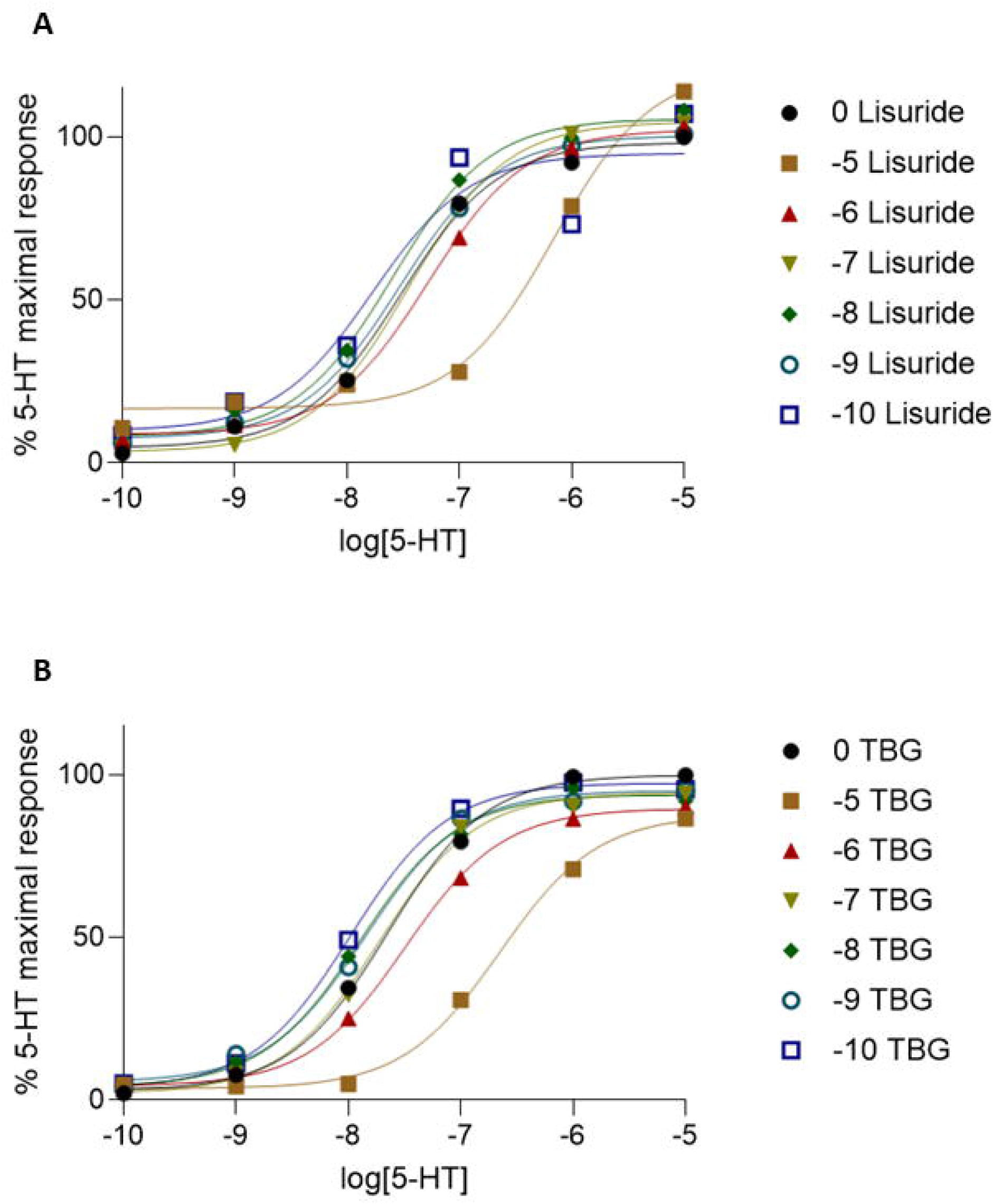
Effect of the non-psychedelic drugs lisuride (A) and TBG (B) on cytosolic Ca^2+^ in C6 cells expressing the rat 5-HT_2A_ receptor in the presence of 5-HT (10 μM). Each point is the mean ± SEM value of triplicates.

Overall, the data from the C6 cells confirmed that the non-psychedelic drugs lisuride and TBG were very low efficacy 5-HT_2A_ receptor agonists, as observed in the SH-SY5Y cells.

## DISCUSSION

It is unknown why some 5-HT_2A_ receptor agonists are psychedelic and others are not, with both biased agonism and partial agonism being plausible explanations (see Introduction). The present study characterised the 5-HT_2A_ receptor signalling properties of a chemically diverse panel of psychedelics (psilocin, 5-MeO-DMT, LSD, mescaline, 25B-NBOMe, DOI) and non-psychedelics (lisuride, TBG), with a focus on G_q_ (cytosolic Ca^2+^, IP_1_ accumulation) versus β-arrestin2 signalling. Key findings were: (i) the psychedelics were 5-HT_2A_ receptor agonists in models of both G_q_ and β-arrestin2 signalling, with these drugs typically being unbiased and having with lower efficacy than 5-HT, (ii) the non-psychedelics, lisuride and TBG were indistinguishable from psychedelics in terms of their biased agonist properties but both exhibited the lowest 5-HT_2A_ receptor signalling efficacy of all drugs tested, and (iii) whilst there was no clear correlation between chemical structure and signalling bias or efficacy, it is noteworthy that the two ergolines, lisuride and LSD, showed evidence of G_q_ signalling bias.

### 5-HT2A receptor signalling bias did not discriminate between psychedelic and non-psychedelic drugs

A potential explanation for why some 5-HT_2A_ receptor agonists are psychedelic and not others is biased agonism - that is, preference for one 5-HT_2A_ receptor signalling pathway versus another. Here, all the psychedelics and non-psychedelics tested activated both 5-HT_2A_ receptor-mediated G_q_ and β-arrestin2 signalling. More importantly, signalling bias did not discriminate between psychedelics and non-psychedelics. Thus, none of the psychedelics showed significant G_q_ versus β-arrestin2 signalling bias (although modest bias was observed for LSD, see below) and of the non-psychedelics, lisuride showed evidence of bias towards G_q_ signalling whereas TBG did not.

Our principal model of G_q_ and β-arrestin2 signalling bias used the human 5-HT_2A_ receptor and measurement of IP_1_ accumulation and β-arrestin2 recruitment combined with a scatter plot of Δlog(E_max_/EC_50_) values for each agonist and in each signalling pathway^44^. A strength of this model is that the use of a reference agonist (here 5-HT) in each assay accounts for assays with different receptor reserves and cell backgrounds as well as any differences in signal amplification. The current study also generated a scatter plot of Δlog(E_max_/EC_50_) values for Ca^2+^ versus β-arrestin2 signalling and this also did not discriminate between the psychedelic and non-psychedelic drugs. However, it should be noted that the latter plot is limited to the extent that Ca^2+^ was measured under non-equilibrium conditions whereas β-arrestin2 measurements were performed at equilibrium (see below).

There are currently few studies of 5-HT_2A_ receptor signalling bias of psychedelic versus non-psychedelic drugs. Our data showing that psilocin and other psychedelics lack bias is in accord with a very recent study also reporting that psilocin, LSD, DOI and 5-MeO-DMT have similar activity at human 5-HT_2A_ receptor-mediated G and β-arrestin2 signalling^34^ although signalling bias was not quantified using Δlog(E /EC) values. Interestingly, an earlier study of human 5-HT_2A_ receptor signalling via IP_1_ and arachidonic acid pathways (downstream of the PLA_2_ pathway) by Berg *et al* reported that DOI, LSD and lisuride showed signalling bias towards the AA pathway^26^. It is possible that psychedelic and non-psychedelics could be discriminated by 5-HT_2A_ receptor signalling via pathways other than G_q_ versus β-arrestin2. However, the finding by Berg *et al* that lisuride showed similar bias to LSD and DOI with regards to IP_1_ versus AA pathways argues against this^26^.

### Non-psychedelic drugs had low efficacy 5-HT_2A_ receptor signalling compared to psychedelic drugs

A general feature of the drugs tested here is that they had lower efficacy compared to 5-HT itself in both G_q_ and β-arrestin2 signalling and were thereby partial 5-HT_2A_ receptor agonists. This finding was robust and consistent across two cell systems (human neuroblastoma SH-SY5Y, rat C6 glioma). Interestingly, the non-psychedelics lisuride and TBG consistently displayed the lowest efficacies of all agonists tested in both cell systems. This finding is in accordance with a recent study by Cao *et al*^16^ which reported that compared to psychedelic drugs, three putative non-psychedelic 5-HT_2A_ receptor agonists (lisuride, IHCH-7079, and IHCH-7086) each exhibited low efficacy in both G_q_ and β-arrestin2 signalling pathways mediated by human 5-HT_2A_ receptors. Similarly, other recent papers have reported that the substituted phenethylamine Ariadne^32^ and ergoline 2-Br-LSD^33^, which are both putative non-psychedelics, show low efficacy in human 5-HT receptor coupled G_q_ and β-arrestin2 signalling pathways compared to psychedelic drugs of similar chemical structure (DOM and LSD, respectively). These findings taken together with the current data suggest that a defining feature of non-psychedelics that differentiates them from psychedelics, is their very low efficacy at 5-HT_2A_ receptors.

Recent evidence suggests that a certain level of efficacy is required for a 5-HT_2A_ receptor agonist to elicit a head-twitch response^34^, a commonly accepted surrogate marker of the psychedelic effect. Specifically, in a study of the 5-HT_2A_ receptor signalling efficacy of 14 phenethylamines, those drugs with low signalling efficacy lacked ability to evoke a head-twitch response. Low efficacy 5-HT_2A_ receptor agonists are apparently also capable of evoking effects claimed to be similar to known antidepressants in preclinical models. For example, low efficacy 5-HT_2A_ receptor agonists such as lisuride, TBG and IHCH-7086 were reported to have such effects in behavioural models and also exhibit increases molecular and cellular markers of plasticity^16,18,33^. It is currently unknown whether this capacity of low efficacy 5-HT_2A_ receptor agonists is of functional relevance in humans.

Partial agonism rather than biased agonism has also been proposed to explain why certain µ-opioid receptor agonists elicit weak respiratory depressant effects (eg. oliceridine) compared to others (eg. morphine and fentanyl)^45–48^. Thus, drugs with weak respiratory depressant effects such as oliceridine had low µ-opioid receptor mediated G_i_ signalling efficacy without significant G_i_ and β-arrestin2 signalling bias^45^. Similarly, it has been proposed that partial agonism explains the actions of behaviourally-selective benzodiazepines (notwithstanding the alternative explanation of GABA_A_ subunit selectivity). In this case, low efficacy drugs such as bretazenil show reduced sedative effects while maintaining anxiolytic effects^35–37^.

### 5-HT2A receptor signalling bias of ergolines

Analysis of G_q_ (IP_1_) versus β-arrestin2 signalling bias revealed some evidence that the ergolines LSD and lisuride had signalling bias properties. Specifically, LSD showed a modest bias towards G_q_ (IP_1_) over β-arrestin2 signalling while lisuride showed a stronger bias in this regard. However, comparison of the data from the IP_1_ and Ca^2+^ assays also revealed that both ergolines exhibited bias towards IP_1_, which should be measuring the same G_q_ signalling pathway as the Ca^2+^ readout. These data suggests that in some cases, measurement of signalling bias may be influenced by assay format.

Here, both IP_1_ and β-arrestin2 measurements were made under conditions in which agonists had likely reached equilibrium with the receptor, and measures were taken as an accumulation of response. Therefore, it is reasonable to conclude that comparison of these two assays provides a more accurate measure of signalling bias. On the other hand, the Ca^2+^ readout was obtained immediately after agonist addition when the drug and receptor were likely under non-equilibrium conditions. When comparisons of agonist activity are made under equilibrium and non-equilibrium conditions, kinetic differences in agonist on- and off-rates as well as receptor residency times can influence measures of agonist efficacy and potency^49,50^, and thereby potentially lead to inaccurate measures of signalling bias.

Our observation that lisuride and LSD had increased 5-HT_2A_ receptor signalling activity in the equilibrium IP_1_ assay versus the non-equilibrium Ca^2+^ assay agrees with previous findings with other ergolines at the 5-HT_2B_ receptor^51,52^, which has high structural homology to the 5-HT_2A_ receptor. Interestingly, it is reported that LSD has a unique 5-HT_2B_ receptor binding mode in which a molecular ‘lid’ hinders drug on- and off-rates and prolongs residency times^38,40^. Presumably this also applies to lisuride. Thus, the Ca^2+^ assay may underestimate the activity of LSD and lisuride due to these drugs not having reached equilibrium with the receptor leading to an apparent IP_1_ versus Ca^2+^ bias. On the other hand, the finding that LSD and lisuride have a G_q_ signalling bias using the IP_1_ and β-arrestin2 assays may be a more accurate finding because both assay conditions were at equilibrium.

Thus, our findings with LSD and lisuride support the contention that the kinetics and equilibrium state of signalling assays is an important consideration when measuring signalling bias parameters.

### Role of 5-HT_2A_ receptor signalling pathways in mediating behavioural effects

Some critical level of 5-HT_2A_ receptor signalling efficacy is required to elicit psychedelic effects in that, as noted above, low efficacy 5-HT_2A_ receptor agonists lack an ability to elicit head-twitches in mice. In the current study there was no clear pattern of 5-HT_2A_ receptor signalling bias from the psychedelic and non-psychedelic drugs tested to inform on the likely pathways mediating the psychedelic effects. Moreover, the literature is currently unclear on this point. Some evidence suggests a role for G_q_ signalling in the head-twitch response. For example, inhibitors of inositol monophosphatase, a key enzyme in the G_q_ signalling pathway, reduced head-twitches induced by DOI and psilocin^53^. Also, G_q_ but not β-arrestin2 signalling efficacy was correlated with propensity to evoke head twitches^34^. However, data generated using β-arrestin2 knockout mice are inconsistent. Whilst one study reported that LSD-induced head-twitches were attenuated in this mouse^54^, other studies found that DOI-induced head-twitches were not^55,56^.

There are similarly conflicting findings regarding how 5-HT_2A_ receptor signalling pathways may mediate different behavioural and neuroplastic effects. Thus, inositol monophosphatase inhibitors were reported to reduce DOI-evoked expression of markers of neural plasticity ^53^, suggesting a role for G_q_ signalling. Accordingly, a phospholipase C inhibitor prevented the increase in plasticity genes in cultured mouse cortical neurons exposed to LSD and lisuride^13^. On the other hand, another study reported that several β-arrestin2-biased 5-HT_2A_ receptor agonists attenuated acute restraint stress-induced freezing behaviour in tail suspension and forced swim tests in mice^16^. A further complication is recent evidence that some behavioural and neuroplastic effects of psychedelic drugs are not mediated by 5-HT_2A_ receptors alone but that the neurotrophic factor receptor TrkB may also play a role^57^.

A final point is that whilst the 5-HT_2A_ receptor signalling pathways underlying the behavioural effects are uncertain, other factors add further uncertainty. In particular, most (if not all) psychedelic and non-psychedelic drugs are not selective 5-HT_2A_ receptor agonists and exhibit affinity for other 5-HT receptors and receptors for other neurotransmitters. For example, in addition to having agonist activity at the 5-HT_2A_ receptor psilocin also exhibits agonist activity at 5-HT_2B/C_ and 5-HT_1A_ receptors, and there is evidence that 5-HT_2C_ receptor activity opposes 5-HT_2A_-mediated effects^58,59^. Thus, the polypharmacology of 5-HT_2A_ agonists likely plays on the behavioural effects of these drugs.

## Conclusion

The present data suggests that the psychedelic drugs tested are not biased towards either 5-HT_2A_ receptor-mediated G_q_ or β-arrestin2 signalling pathways. The non-psychedelic 5-HT_2A_ receptor agonists lisuride and TBG also did not have a consistent bias. Rather a feature of the latter drugs was their low 5-HT_2A_ receptor efficacy in both G_q_ and β-arrestin2 signalling pathways. This finding combined with other recent studies reporting low efficacy of other non-psychedelic 5-HT_2A_ receptor agonists, suggests that low efficacy rather than signalling bias plays a key role in their lack of psychedelic effect.

## Acknowledgements

This study was supported by a MRC iCASE studentship (AI) with Compass Pathways plc.

The data that support the findings of this study are available from the corresponding author upon reasonable request.

## References

1. Ross, S. et al. Rapid and sustained symptom reduction following psilocybin treatment for anxiety and depression in patients with life-threatening cancer: a randomized controlled trial. J. Psychopharmacol. 30, 1165–1180 (2016).

2. Griffiths, R. R. et al. Psilocybin produces substantial and sustained decreases in depression and anxiety in patients with life-threatening cancer: A randomized double-blind trial. J. Psychopharmacol. 30, 1181–1197 (2016).

3. Carhart-Harris, R. et al. Trial of Psilocybin versus Escitalopram for Depression. N. Engl. J. Med. 384, 1402–1411 (2021).

4. Holze, F., Gasser, P., Müller, F., Dolder, P. C. & Liechti, M. E. Lysergic Acid Diethylamide-Assisted Therapy in Patients With Anxiety With and Without a Life-Threatening Illness: A Randomized, Double-Blind, Placebo-Controlled Phase II Study. Biol. Psychiatry 93, 215–223 (2023).

5. Reckweg, J. T. et al. A phase 1/2 trial to assess safety and efficacy of a vaporized 5-methoxy-N,N-dimethyltryptamine formulation (GH001) in patients with treatment-resistant depression. Front. psychiatry 14, 1133414 (2023).

6. Madsen, M. K. et al. Psychedelic effects of psilocybin correlate with serotonin 2A receptor occupancy and plasma psilocin levels. Neuropsychopharmacol. Off. Publ. Am. Coll. Neuropsychopharmacol. 36, 45–73 (2018).

7. Sharp, T. & Barnes, N. M. Central 5-HT receptors and their function; present and future. Neuropharmacology 177, 108155 (2020).

8. Vollenweider, F. X., Vollenweider-Scherpenhuyzen, M. F., Bäbler, A., Vogel, H. & Hell, D. Psilocybin induces schizophrenia-like psychosis in humans via a serotonin-2 agonist action. Neuroreport 9, 3897–3902 (1998).

9. Valle, M. et al. Inhibition of alpha oscillations through serotonin-2A receptor activation underlies the visual effects of ayahuasca in humans. Eur. Neuropsychopharmacol. J. Eur. Coll. Neuropsychopharmacol. 26, 1161–1175 (2016).

10. Preller, K. H. et al. The Fabric of Meaning and Subjective Effects in LSD-Induced States Depend on Serotonin 2A Receptor Activation. Curr. Biol. 27, 451–457 (2017).

11. Holze, F. et al. Acute dose-dependent effects of lysergic acid diethylamide in a double-blind placebo-controlled study in healthy subjects. Neuropsychopharmacol. Off. Publ. Am. Coll. Neuropsychopharmacol. 46, 537–544 (2021).

12. Becker, A. M. et al. Ketanserin Reverses the Acute Response to LSD in a Randomized, Double-Blind, Placebo-Controlled, Crossover Study in Healthy Participants. Int. J. Neuropsychopharmacol. 26, 97– 106 (2023).

13. González-Maeso, J. et al. Hallucinogens recruit specific cortical 5-HT(2A) receptor-mediated signaling pathways to affect behavior. Neuron 53, 439–452 (2007).

14. Jennings, K. A., Sheward, W. J., Harmar, A. J. & Sharp, T. Evidence that genetic variation in 5-HT transporter expression is linked to changes in 5-HT2A receptor function. Neuropharmacology 54, 776–783 (2008).

15. Halberstadt, A. L., Chatha, M., Klein, A. K., Wallach, J. & Brandt, S. D. Correlation between the potency of hallucinogens in the mouse head-twitch response assay and their behavioral and subjective effects in other species. Neuropharmacology 167, (2020).

16. Dongmei, C. et al. Structure-based discovery of nonhallucinogenic psychedelic analogs. Science (80-.). 375, 403–411 (2022).

17. Dong, C. et al. Psychedelic-inspired drug discovery using an engineered biosensor. Cell 184, 2779–2792.e18 (2021).

18. Cameron, L. P. et al. A non-hallucinogenic psychedelic analogue with therapeutic potential. Nature (2020) doi:10.1038/s41586-020-3008-z.

19. Claus, J. J. et al. Lisuride treatment of Alzheimer’s disease. A preliminary placebo-controlled clinical trial of safety and therapeutic efficacy. Clin. Neuropharmacol. 21, 190–195 (1998).

20. Herrmann, W. M., Horowski, R., Dannehl, K., Kramer, U. & Lurati, K. Clinical effectiveness of lisuride hydrogen maleate: a double-blind trial versus methysergide. Headache 17, 54–60 (1977).

21. Schmidt, L. G., Kuhn, S., Smolka, M., Schmidt, K. & Rommelspacher, H. Lisuride, a dopamine D2 receptor agonist, and anticraving drug expectancy as modifiers of relapse in alcohol dependence. Prog. Neuropsychopharmacol. Biol. Psychiatry 26, 209–217 (2002).

22. Urban, J. D. et al. Functional selectivity and classical concepts of quantitative pharmacology. J. Pharmacol. Exp. Ther. 320, 1–13 (2007).

23. Kenakin, T. Agonist-receptor efficacy. II. Agonist trafficking of receptor signals. Trends Pharmacol. Sci. 16, 232–238 (1995).

24. Reiter, E., Ahn, S., Shukla, A. K. & Lefkowitz, R. J. Molecular mechanism of β-arrestin-biased agonism at seven-transmembrane receptors. Annu. Rev. Pharmacol. Toxicol. 52, 179–197 (2012).

25. Kurrasch-Orbaugh, D. M., Watts, V. J., Barker, E. L. & Nichols, D. E. Serotonin 5-hydroxytryptamine 2A receptor-coupled phospholipase C and phospholipase A2 signaling pathways have different receptor reserves. J. Pharmacol. Exp. Ther. 304, 229–237 (2003).

26. Berg, K. A. et al. Effector pathway-dependent relative efficacy at serotonin type 2A and 2C receptors: evidence for agonist-directed trafficking of receptor stimulus. Mol. Pharmacol. 54, 94– 104 (1998).

27. Xia, Z., Gray, J. A., Compton-Toth, B. A. & Roth, B. L. A direct interaction of PSD-95 with 5-HT2A serotonin receptors regulates receptor trafficking and signal transduction. J. Biol. Chem. 278, 21901–21908 (2003).

28. Xia, Z., Hufeisen, S. J., Gray, J. A. & Roth, B. L. The PDZ-binding domain is essential for the dendritic targeting of 5-HT2A serotonin receptors in cortical pyramidal neurons in vitro. Neuroscience 122, 907–920 (2003).

29. Pottie, E. et al. Structure-Activity Assessment and In-Depth Analysis of Biased Agonism in a Set of Phenylalkylamine 5-HT(2A) Receptor Agonists. ACS Chem. Neurosci. 14, 2727–2742 (2023).

30. Bohn, L. M. et al. Enhanced morphine analgesia in mice lacking beta-arrestin 2. Science 286, 2495– 2498 (1999).

31. Bohn, L. M., Gainetdinov, R. R., Lin, F. T., Lefkowitz, R. J. & Caron, M. G. Mu-opioid receptor desensitization by beta-arrestin-2 determines morphine tolerance but not dependence. Nature 408, 720–723 (2000).

32. Cunningham, M. J. et al. Pharmacological Mechanism of the Non-hallucinogenic 5-HT(2A) Agonist Ariadne and Analogs. ACS Chem. Neurosci. 14, 119–135 (2023).

33. Lewis, V. et al. A non-hallucinogenic LSD analog with therapeutic potential for mood disorders. Cell Rep. 42, 112203 (2023).

34. Wallach, J. et al. Identification of 5-HT(2A) Receptor Signaling Pathways Responsible for Psychedelic Potential. bioRxiv,: the preprint server for biology at 10.1101/2023.07.29.551106 (2023).

35. Jackson, H. C. Benzodiazepine partial agonists. J. Psychopharmacol. 7, 101–103 (1993).

36. Haefely, W. et al. Partial agonists of benzodiazepine receptors for the treatment of epilepsy, sleep, and anxiety disorders. Adv. Biochem. Psychopharmacol. 47, 379–394 (1992).

37. Haefely, W., Martin, J. R. & Schoch, P. Novel anxiolytics that act as partial agonists at benzodiazepine receptors. Trends Pharmacol. Sci. 11, 452–456 (1990).

38. McCorvy, J. D. et al. Structural determinants of 5-HT(2B) receptor activation and biased agonism. Nat. Struct. Mol. Biol. 25, 787–796 (2018).

39. Wacker, D. et al. Structural features for functional selectivity at serotonin receptors. Science 340, 615–619 (2013).

40. Kim, K. et al. Article Structure of a Hallucinogen-Activated Gq-Coupled 5-HT 2A Serotonin Receptor ll Article Structure of a Hallucinogen-Activated Gq-Coupled 5-HT 2A Serotonin Receptor. Cell 182, 1574–1588.e19 (2020).

41. Newton, R. A. et al. Characterisation of human 5-hydroxytryptamine2A and 5-hydroxytryptamine2C receptors expressed in the human neuroblastoma cell line SH-SY5Y: comparative stimulation by hallucinogenic drugs. J. Neurochem. 67, 2521–2531 (1996).

42. Meller, R., Harrison, P. J. & Sharp, T. Studies on the role of calcium in the 5-HT-stimulated release of glutamate from C6 glioma cells. Eur. J. Pharmacol. 445, 13–19 (2002).

43. Meller, R., Harrison, P. J., Elliott, J. M. & Sharp, T. In vitro evidence that 5-hydroxytryptamine increases efflux of glial glutamate via 5-HT(2A) receptor activation. J. Neurosci. Res. 67, 399–405 (2002).

44. Kenakin, T., Watson, C., Muniz-Medina, V., Christopoulos, A. & Novick, S. A simple method for quantifying functional selectivity and agonist bias. ACS Chem. Neurosci. 3, 193–203 (2012).

45. Gillis, A. et al. Low intrinsic efficacy for G protein activation can explain the improved side effect profiles of new opioid agonists. Sci. Signal. 13, (2020).

46. Rivero, G. et al. Endomorphin-2: a biased agonist at the μ-opioid receptor. Mol. Pharmacol. 82, 178– 188 (2012).

47. McPherson, J. et al. μ-opioid receptors: correlation of agonist efficacy for signalling with ability to activate internalization. Mol. Pharmacol. 78, 756–766 (2010).

48. Yudin, Y. & Rohacs, T. The G-protein-biased agents PZM21 and TRV130 are partial agonists of μ-opioid receptor-mediated signalling to ion channels. Br. J. Pharmacol. 176, 3110–3125 (2019).

49. Kenakin, T. P. The classification of drugs and drug receptors in isolated tissues. Pharmacol. Rev. 36, 165–222 (1984).

50. Finlay, D. B., Duffull, S. B. & Glass, M. 100 years of modelling ligand–receptor binding and response: A focus on GPCRs. Br. J. Pharmacol. 177, 1472–1484 (2020).

51. Unett, D. J. et al. Kinetics of 5-HT2B Receptor Signaling: Profound Agonist-Dependent Effects on Signaling Onset and Duration. J. Pharmacol. Exp. Ther. 347, 645–659 (2013).

52. Bdioui, S. et al. Equilibrium assays are required to accurately characterize the activity profiles of drugs modulating Gq-protein-coupled receptors. Mol. Pharmacol. 94, 992–1006 (2018).

53. Antoniadou, I. et al. Ebselen has lithium-like effects on central 5-HT(2A) receptor function. Br. J. Pharmacol. 175, 2599–2610 (2018).

54. Rodriguiz, R. M. et al. LSD-stimulated behaviors in mice require β-arrestin 2 but not β-arrestin 1. Sci. Rep. 11, 17690 (2021).

55. de la Fuente Revenga, M., et al. Tolerance and Cross-Tolerance among Psychedelic and Nonpsychedelic 5-HT(2A) Receptor Agonists in Mice. ACS Chem. Neurosci. 13, 2436–2448 (2022).

56. Schmid, C. L., Raehal, K. M. & Bohn, L. M. Agonist-directed signaling of the serotonin 2A receptor depends on beta-arrestin-2 interactions in vivo. Proc. Natl. Acad. Sci. U. S. A. 105, 1079–1084 (2008).

57. Moliner, R. et al. Psychedelics promote plasticity by directly binding to BDNF receptor TrkB. Nat. Neurosci. 26, 1032–1041 (2023).

58. Halberstadt, A. L. et al. 5-HT(2A) and 5-HT(2C) receptors exert opposing effects on locomotor activity in mice. Neuropsychopharmacol. Off. Publ. Am. Coll. Neuropsychopharmacol. 34, 1958– 1967 (2009).

59. Fantegrossi, W. E. et al. Interaction of 5-HT2A and 5-HT2C receptors in R(-)-2,5-dimethoxy-4-iodoamphetamine-elicited head twitch behavior in mice. J. Pharmacol. Exp. Ther. 335, 728–734 (2010).

