## Supplementary Data for "Increased 5-HT_2A_ receptor signalling efficacy differentiates serotonergic psychedelics from non-psychedelics"

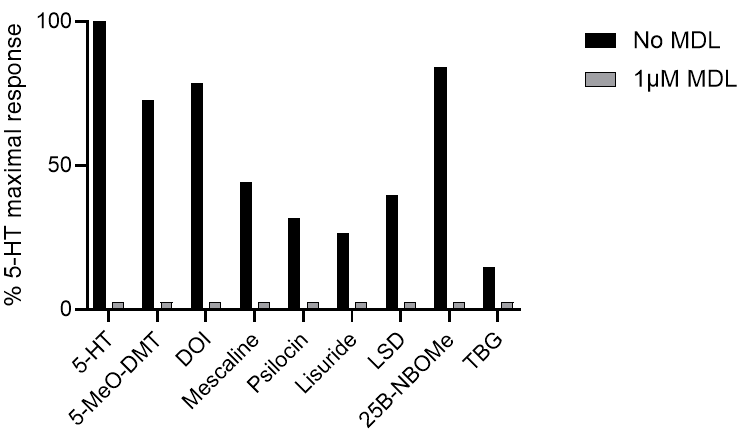

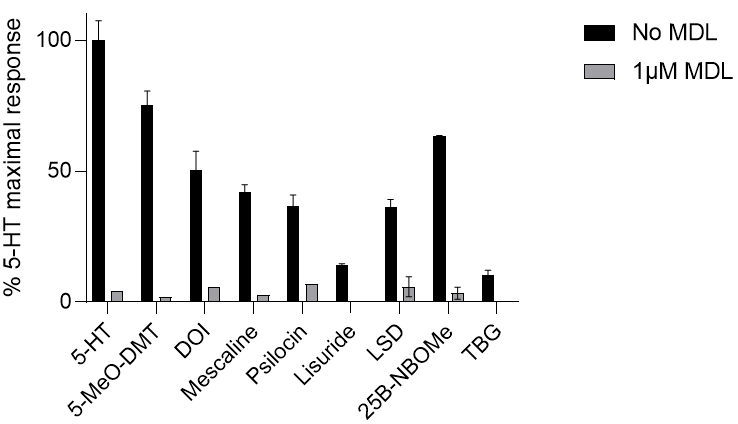

**B**

**A**

**Figure S1.** Effect of psychedelic and non-psychedelic drugs (10 µM) on A) Ca^2+^ and B) β-arrestin2 responses in SH-SY5Y cells in the presence or absence of the selective 5-HT_2A_ receptor antagonist MDL-100907 (1 µM). Each point represents a mean ± SEM value. Responses are relative to 5-HT (10 μM). In this experiment responses obtained in the presence of MDL-100907 were undetectable, so corresponding columns are for illustration purposes only.

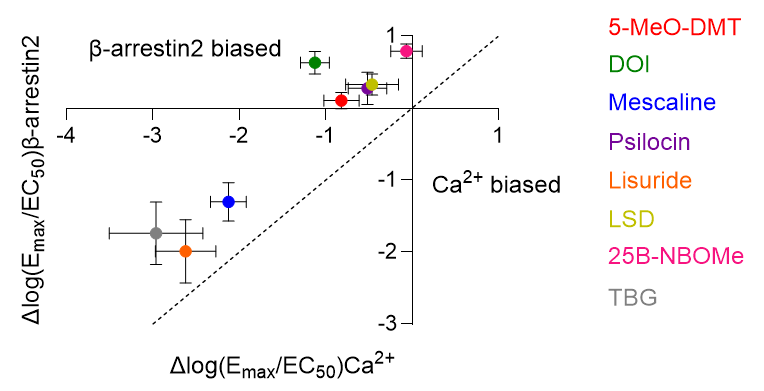

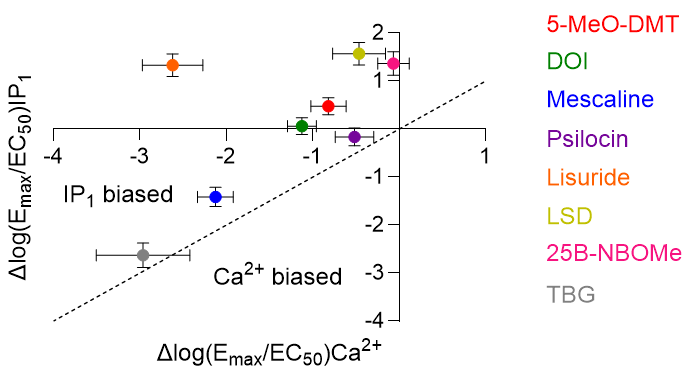

**B**

**A**

**Figure S2 |** Scatter plots comparing the activity (Δlog(E_max_/EC_50_) values) of psychedelic and non-psychedelic drugs on A) 5-HT_2A_ receptor-mediated IP_1_ and Ca^2+^ signalling pathways and B) 5-HT_2A_ receptor-mediated β-arrestin and Ca^2+^ signalling pathways in SH-SY5Y cells. In this plot, the more positive the x or y value, the greater activity in a particular pathway. The further a drug deviates from the line of unity, the more biased the agonist. Each point represents a mean ± SEM value.

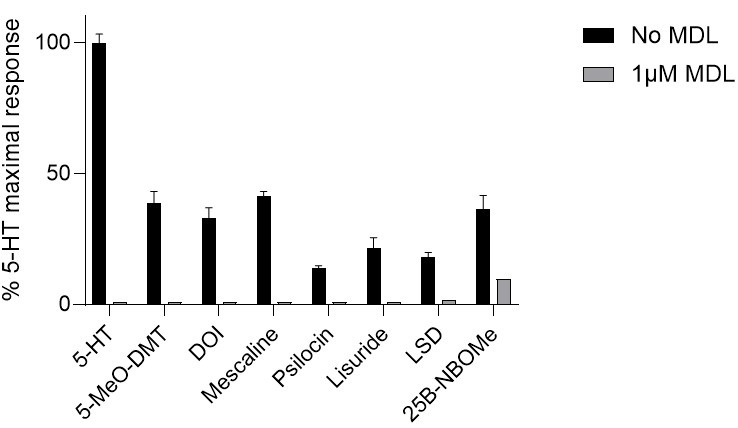

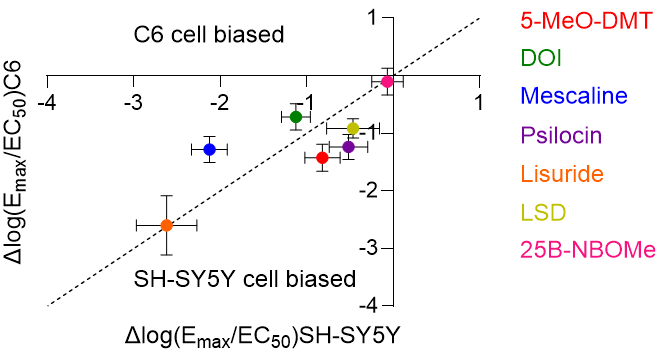

**B**

**A**

**Figure S3.** A) Scatter plot comparing the activity (Δlog(E_max_/EC_50_) values) of psychedelic and non-psychedelic drugs on Ca^2+^ signalling in SH-SY5Y versus C6 cells. B) Effect of psychedelic and non-psychedelic drugs (10 µM) on Ca^2+^ responses in C6 cells in the presence or absence of the selective 5-HT_2A_ receptor antagonist MDL-100907 (1 µM). Each point represents a mean ± SEM value. In this experiment most responses obtained in the presence of MDL-100907 were undetectable, so corresponding columns are for illustration purposes only.

**Supplementary Table 1**

Schild plot slopes, linear regression and pA_2_ values for psychedelic and non-psychedelic drugs calculated from Ca^2+^ responses in the presence of 5-HT (10 μM) in C6 cells. * = pA_2_ value calculated despite a Schild plot slope significantly different from 1.

| **Agonist** | **Schild plot slope** | **Linear regression R^2^** | **pA_2_** |
| --- | --- | --- | --- |
| **5-MeO-DMT** | 1.00 | 0.982 | -6.31 |
| **DOI** | 0.362 | 0.832 | -9.37* |
| **Mescaline** | 0.927 | 0.883 | -6.76 |
| **Psilocin** | 1.33 | 0.969 | -6.66 |
| **Lisuride** | 1.03 | 0.947 | -6.17 |
| **LSD** | 0.900 | 0.993 | -6.68 |
| **25B-NBOMe** | 0.474 | 0.756 | -8.53* |
| **TBG** | 1.32 | 1.00 | -5.72 |
